# Variants in *RABL2A* causing male infertility and ciliopathy

**DOI:** 10.1101/2020.08.11.246611

**Authors:** Xinbao Ding, Robert Fragoza, Priti Singh, Shu Zhang, Haiyuan Yu, John C Schimenti

## Abstract

**Purpose:** Approximately 7% of men suffer from infertility worldwide and sperm abnormalities are the major cause. Though genetic defects are thought to underlie a substantial fraction of all male infertility cases, the actual molecular bases are usually undetermined. Because the consequences of most genetic variants in populations are unknown, this complicates genetic diagnosis even after genome sequencing of patients. Some patients with ciliopathies, including primary ciliary dyskinesia (PCD) and Bardet-Biedl syndrome (BBS), also suffer from infertility because sperm flagella, which share several characteristics with cilia, are also affected in these patients.

**Methods:** To identify infertility-causing genetic variants in human populations, we used *in silico* predictions to identify potentially deleterious SNP (single nucleotide polymorphism) alleles of *RABL2A*, a gene essential for normal cilia and flagella function. Candidate variants were assayed for protein stability *in vitro*, and the destabilizing variants were modeled in mice using CRISPR/Cas9-mediated genome editing. The resulting mice were characterized phenotypically for reproductive and developmental defects.

**Results:** Two of the SNP alleles, *Rabl2*^*L119F*^ (rs80006029) and *Rabl2*^*V158F*^ (rs200121688), destabilized the protein. Mice bearing these alleles exhibited PCD- and BBS-associated disorders including male infertility, early growth retardation, excessive weight gain in adulthood, heterotaxia, pre-axial polydactyly, neural tube defects (NTD) and hydrocephalus.

**Conclusion:** Our study identified and validated pathogenicity of two variants causing ciliopathies and male infertility in human populations, and identified phenotypes not previously described for null alleles of *Rabl2*.

## INTRODUCTION

Infertility affects approximately 7% of the male population worldwide ^1^ and often manifests qualitative and/or quantitative defects in sperm parameters. Sperm tail (flagellum) defects can cause teratospermia (abnormal morphology), asthenospermia (reduced sperm motility), azoospermia and oligospermia (no and low sperm count, respectively), thus contributing to main categories of male infertility. Asthenozoospermia is often observed in patients suffering from primary ciliary dyskinesia (PCD) (OMIM no. 244400), a group of autosomal-recessive disorders caused by motile cilia dysfunction. Cilia are categorized into two types, motile cilia/flagella and non-motile cilia (also called primary cilia). The former is responsible for propelling cells or generating fluid flow over ciliated cells, whereas the latter functions as cellular antennae by sensing extracellular stimuli, and receiving and transducing developmental signals. Defects in cilia result in a variety of congenital disorders, such as PCD, Bardet-Biedl syndrome (BBS), Joubert syndrome (JBTS) and Meckel-Grüber syndrome (MKS), which are collectively called the ciliopathies ^2^. PCD results in recurrent respiratory infections, male infertility, randomization of left-right asymmetry and hydrocephalus. BBS is characterized by features including cognitive impairment, obesity, retinal degeneration, anosmia, neural tube defects (NTDs), cystic kidneys and polydactyly. MKS and JBTS are characterized by severe central nervous system (CNS) disorders.

Rab-like protein 2 (RABL2) is a small GTPase member of the Ras superfamily and is essential for proper function of flagella and cilia. In flagella, RABL2 localizes to the mid-piece region and can deliver a specific set of effector proteins including key members of the glycolytic pathway into the growing sperm tails by binding Intraflagellar Transport-B (IFT-B; a large of proteins) and guanosine triphosphate (GTP) ^3^. In cilia, RABL2 can be recruited by CEP19 (centrosomal protein 19) to the mother centriole, bind GTP and the IFT-B complex, triggering its entry into the cilium ^4^. The CEP19-RABL2-IFT pathway was proposed as a molecular mechanism for directing ciliary traffic, needed for cilium formation. A mouse model called “*Mot*” was isolated in an ENU (N-ethyl-N-nitrosourea) mutagenesis screen, which carries a mutation (p.Asp73Gly) in *Rabl2* ^3^. Homozygous mutant *Rabl2*^*Mot*^ mice exhibited qualitatively normal spermatogenesis but were sterile due to the reduced sperm motility and oligospermia ^3^. Besides infertility, *Mot* mice also exhibited adult onset obesity and fatty livers, as well as impaired glucose and lipid metabolism ^5^. In addition, knockout *Rabl2* mice display retinal degeneration and pre-axial polydactyly ^4^, which, together with the obesity and sperm defects, are reminiscent of symptoms observed in patients suffering from PCD and BBS. In humans, there are two highly similar paralogs, *RABL2A* (OMIM no. 605412) and *RABL2B*, in which the extra copy arose after the split between humans and orangutans ^6^. A deletion allele rs57719031 in *RABL2A* has been identified as a potential risk factor for Australian infertility patients ^7^. However, it was not reported whether these patients presented PCD- or BBS-associated symptoms.

It is important to characterize genetic causes of disease to aid in clinical diagnosis and possibly treatment. SNPs are the most prevalent type of genetic variation in both coding and non-coding regions of the genome. Nonsynonymous SNPs (nsSNPs; those which change an amino acid) can play key roles in causing disease by affecting protein structure or function, protein-macromolecules interactions, or post-translational modifications ^8^. However, the effects of most SNPs are unknown, and are referred to as variants of unknown significance (VUS) ^9^. Identifying those that contribute to disease is a major challenge in human genetics. To identify VUS that cause infertility in human populations, we have implemented an association- and linkage-free approach ^10^. This approach involves selecting nsSNPs in genes known to be required for gametogenesis and fertility, identifying those nsSNPs that are predicted computationally to be deleterious, modeling the orthologous amino acid changes in mice using CRISPR/Cas9 genome editing, then assessing phenotypes of the mice ^10^. To this end, several nsSNPs affecting infertility were identified ^10–13^.

Here, we present evidence that two nsSNP alleles of *RABL2A*, rs80006029 and rs200121688, are functionally deleterious. These variants disrupt protein stability, and when modeled in mice, caused PCD and BBS-associated disorders including male infertility, excessive weight, growth retardation, heterotaxia, pre-axial polydactyly, NTDs and hydrocephalus.

## MATERIALS AND METHODS

### Computational identification of deleterious nsSNPs

The possible pathogenic functional effects of nsSNPs on human RABL2A were analyzed using SIFT ^14^ and PolyPhen-2 ^15^. The frequency of variants is as reported in gnomAD version 3.0 (gnomeAD.broadinstitute.org). To predict the functional consequences of the mutations, we modelled p.Ala94Thr, p.Leu119Phe and p.Val158Phe variants into the human RABL2A 3D structure (SWISS-MODEL accession code Q9UBK7), and used the following web servers: I-Mutant ^16^, DynaMut ^17^, SDM ^18^, DeepDDG ^19^, DUST ^20^ and mCSM ^21^. PyMol software was used to visualize the RABL2A 3D structure and protein destabilization.

### Constructing vectors for DUAL-FLUO screen and Western blot

Vector construction and Western blot were performed as previous described ^22^. Briefly, the WT and mutant RABL2A open reading frames were inserted into pDEST-DUAL vector by Gateway LR reactions and then the vectors were transfected into HEK293T cells. Anti-GFP (1:1000, SCBT, sc-9996) and anti-GAPDH (1:3000, Proteintech, 60004-1-Ig) were used for immunoblotting analyses.

### Production of CRISPR/Cas9 edited mice

All animal usage was approved by Cornell University’s Institutional Animal Care and Use Committee, under protocol 2004-0038 to J.C.S.

*Rabl2* mutant mice were generated using CRISPR/Cas9-mediated homologous recombination in zygotes, as described previously ^10^. Design and selection of sgRNAs took into account the parameters of 1) on-target ranking, 2) minimal predicted off-target sites, and 3) the distance to the target site ^23^. The sgRNAs were produced using MEGAshortscript™ T7 Transcription Kit (Ambion; AM1354). sgRNAs and ssODNs are listed in Table S1. Briefly, the sgRNA, ssODN and Cas9 mRNA (25ng/μL, TriLink) were co-injected into zygotes (F1 hybrids between strains FVB/NJ and B6(Cg)-Tyr^c-2J^/J) then transferred into the oviducts of pseudopregnant females. Founders carrying at least one copy of the desired alteration were identified and backcrossed into FVB/NJ. Initial phenotyping was done after one backcross generation and additional phenotyping was done with mice backcrossed two or more generations.

### Genotyping

Mice were genotyped by PCR followed by Sanger sequencing. The PCR primer sequences are listed in Table S1. PCR was performed using EconoTaq and associated PCR reagents (Lucigen) with 4 μL of crude DNA lysate created as described previously ^24^ from ear punch biopsy specimens of 8 to 14-day-old mice. Annealing temperature was 56 °C for *Rabl2*^*LF/LF*^ and *Rabl2*^*-/-*^, 60 °C for *Rabl2*^*VF/VF*^.

### qRT-PCR

Total RNA was extracted from testis by TRIzol (ThermoFisher), and the reverse transcription was performed using a qScript™ cDNA SuperMix (Quantabio, 95048), according to the manufacturer’s instructions. For qRT-PCR, the specificity of PCRs was verified by a single peak according to melt curves. The primer sequences are listed in Table S1. qRT-PCR was performed with a C1000 Touch™ Thermal Cycler (Bio-Rad) amplification system using RT^2^ SYBR^®^ Green qPCR Mastermixes (Qiagen). The relative levels of transcripts were calculated using the 2^-ΔΔCT^ method and the *Rabl2* expression levels were normalized to *Gapdh*.

### Fertility testing

For fertility tests, 2-month-old mice were placed with age-matched FVB/NJ mates. After a period of at least 3 months without offspring, the analyzed individuals were considered infertile.

### Histology and imaging

For the preparation of paraffin blocks, tissues were fixed overnight in Bouin’s solution or 4% PFA (paraformaldehyde) at room temperature, washed in 70% ethanol, then dehydrated and embedded in paraffin. For histological analyses, paraffin sections at 6 μm thick were deparaffinized, then stained with hematoxylin and eosin (H&E). H&E slides were examined on an Olympus BX51 microscope using a 10X objective, and Olympus cellSens standard software. Cropping, color, and contrast adjustments were made with Adobe Photoshop CC 2019, using identical background adjustments for all images. Cauda epididymal sperm tail length was measured following staining with H&E. Forty tails per mouse were measured using Olympus cellSens software.

### Sperm counts

Epididymal sperm counting was performed using an adaptation of a published method ^25^. One cauda epididymis per male was used for each data point.

### Computer-assisted sperm analysis (CASA)

Vas deferens were harvested from adult males, and placed in a puddle of (500 μL) *in vitro* fertilization (IVF) media (Research Vitro Fert K-RVFE-50; Cook Medical, Inc. USA). The semen was squeezed out by the needle and the sperm was allowed to swim out for 10 min at 37 °C. A 150 μL drop of sperm suspension was placed into the same volume of pre-warmed IVF media (37 °C). Then the sperm were moved to a pre-warmed glass slide for motility assessment on an IVOS SpermAnalyzer (Hamilton-Thorne Research, Beverly, Mass.).

### Statistics

All data were expressed as mean ± SEM. Statistical calculations were carried out using a student’s t-test or one-way analysis of variance (ANOVA) followed by the Tukey’s *post hoc* test or Chi-squared test with Statistical Package for the Social Sciences (SPSS) software or within R. Graph generation was performed using R software.

## RESULTS

### Prioritizing potentially deleterious missense variants in *RABL2A*

To identify potentially deleterious segregating mutations in *RABL2A*, we used gnomAD (Genome Aggregation Database), a database containing whole-exome sequencing data for >140,000 individuals ^26^, to identify missense variants in *RABL2A* with a minor allele frequency (MAF) > 0.02%. Six missense variants fulfilled our criteria (Table 1). Since common variants are often presumed to be functionally benign, we further prioritized these six *RABL2A* mutations using SIFT^14^ and PolyPhen-2^15^ and found that p.L119F, p.A94T, and p.V158F scored as deleterious by both algorithms (Table 1). Mutations predicted as being deleterious by variant classifiers may not always have phenotypic consequences ^10,11,13^. Structural analysis suggested that these three mutations destabilize protein folding (Fig 1a and Fig S1). We therefore tested whether these three mutations indeed impacted protein stability through western blot analysis of GFP-tagged versions of these proteins overexpressed in HEK293T cells. The p.L119F and p.V158F variants caused far lower levels of RABL2A protein in comparison to wild-type (WT), while protein levels of A94T remained largely intact (Fig 1b and c). These results suggest that *RABL2A* variants p.L119F and p.V158F are deleterious while p.A94T does not markedly impact protein stability. We therefore selected p.L119F and p.V158F for *in vivo* validation in mice.

**Table 1.** Selection of deleterious nsSNPs.

| **Variant** | **SNP ID** | **MAF** | **SIFT** | **PolyPhen-2** |
| --- | --- | --- | --- | --- |
| **p. Leu119Phe** | **rs80006029** | **0.80%** | **Deleterious** | **Probably Damaging** |
| p. Arg82Gln | rs145167719 | 0.42% | Tolerated | Possibly Damaging |
| p. Glu217Lys | rs142229156 | 0.37% | Tolerated LC* | Benign |
| **p. Ala94Thr** | **rs145909437** | **0.13%** | **Deleterious** | **Probably Damaging** |
| p. Arg120Gln | rs564553154 | 0.035% | Tolerated | Possibly Damaging |
| **p. Val158Phe** | **rs200121688** | **0.023%** | **Deleterious** | **Probably Damaging** |

\* LC: low confidence

**Fig. 1.**
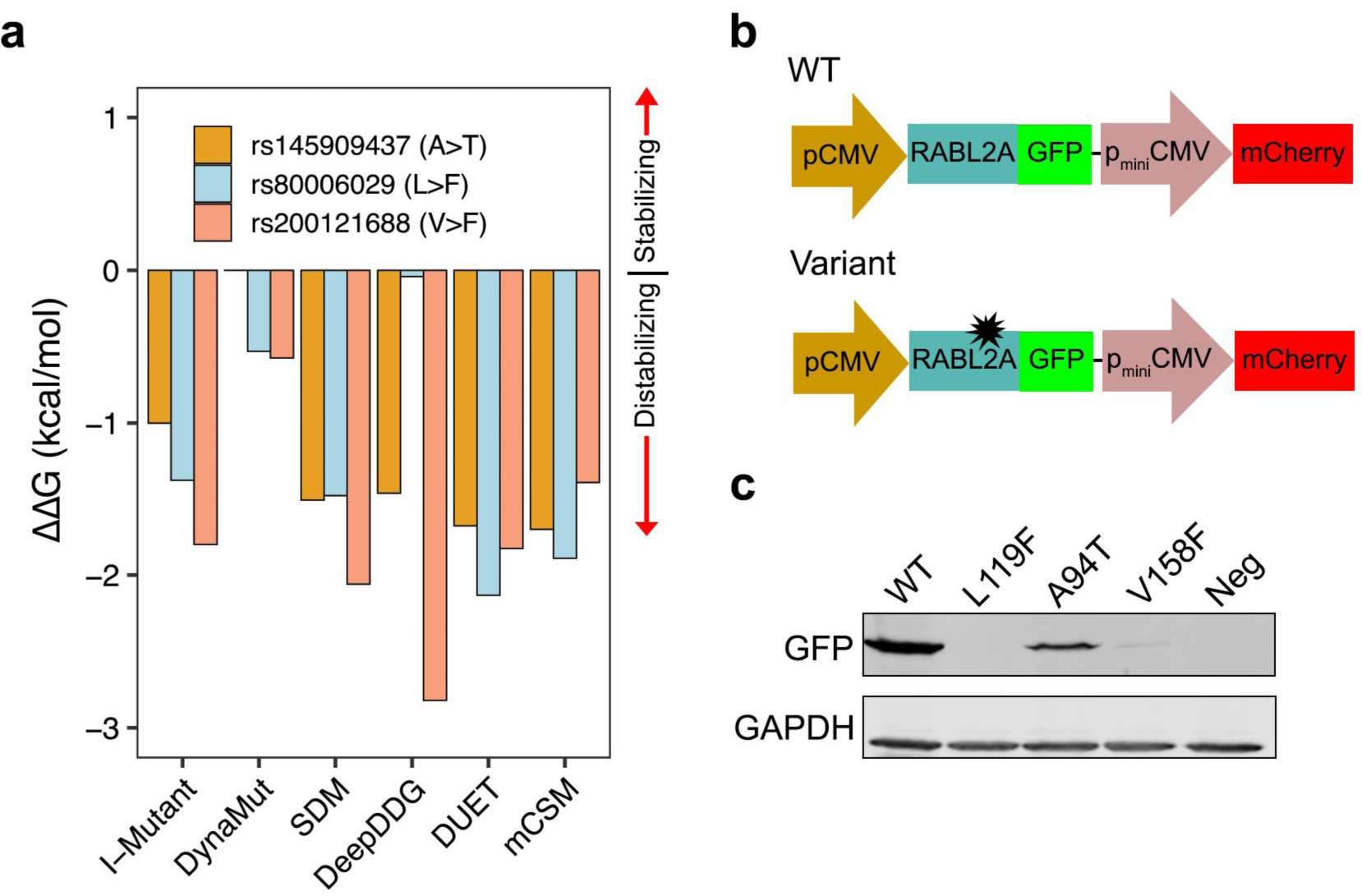
Experimental test the selected nsSNPs. (**a**) Prediction of ΔΔG in p.Ala94Thr, p.Leu119Phe and p.Val158Phe variants by six protein stability prediction algorithms. For I-Mutant, ΔΔG (1 kcal/mol=4,184 J) <-0.5 indicates large decrease of stability, ΔΔG>0.5 indicates large increase of stability and -0.5≤ΔΔG≤0.5 represents neutral stability. For DynaMut, SDM, DeepDDG, DUET and mCSM, positive ΔΔG indicates stabilization by the specified mutation, negative ΔΔG indicates destabilization. (**b**) Schematic of the pDEST-DUAL vector with RABL2A WT and variant ORFs used for protein stability assay. (**c)** Western blots for representative WT and variants detected using α-GFP. α-GAPDH was used as a loading control. Cells transfected with empty plasmid used as negative control.

### *Rabl2* variants cause male infertility in mice

To model the p.L119F and p.V158F conversions encoded by rs80006029 and rs200121688, respectively, equivalent point mutations in mouse *Rabl2* were induced by CRISPR/Cas9 mediated genome editing (Fig 2a). Founder mice with the correct mutation were backcrossed into strain FvB/nJ for two generations. A line with a 1nt frameshift mutation in exon 5 was also established to serve as a presumptive null allele (*Rabl2*^*-/-*^) for comparison. Consistent with a reports of *Rabl2*^*Mot* 3^ and null mice ^4^, *Rabl2*^*V158F/V158F*^ (*Rabl2*^*VF/VF*^), *Rabl2*^*L119F/L119F*^ (*Rabl2*^*LF/LF*^), and *Rabl2*^*-/-*^ mice were all viable (Fig 2b). qRT-PCR analysis indicated that testicular *Rabl2* mRNA levels in the SNP alleles were comparable to WT, but lower in the nulls. (Fig 2c).

**Fig. 2.**
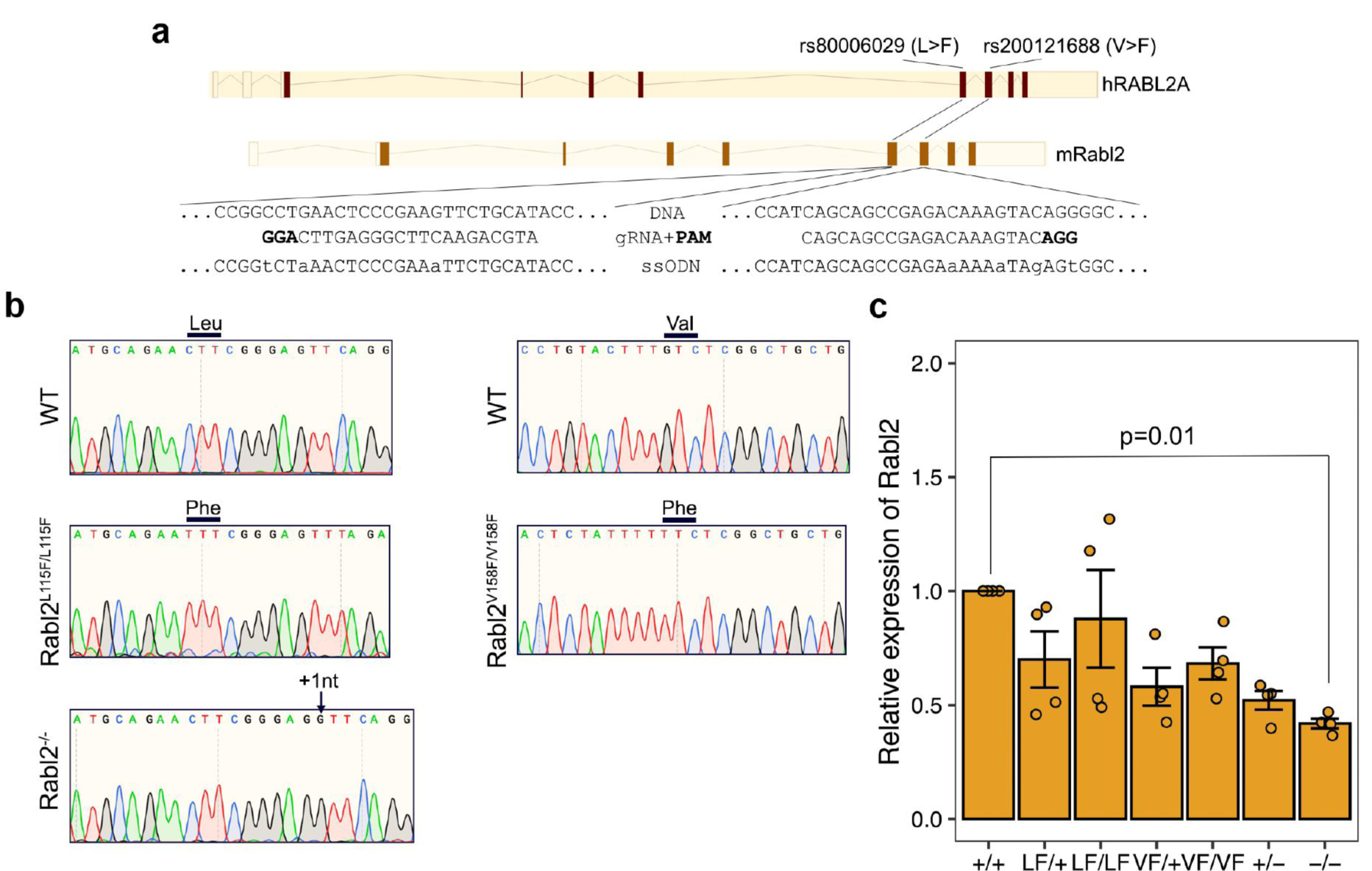
CRISPR/Cas9-mediated generation of Rabl2 mutant mice. (**a**) Diagram of CRISPR-Cas9 genome editing strategy to introduce the p.Leu119Phe and p.Val158Phe amino acid changes into mouse *Rabl2*. The human RABL2A p.Leu119Phe and p.Val158Phe variants encoded by SNP rs80006029 and rs200121688 are located in exons 5 and 6, respectively. (**b**) Representative Sanger sequencing chromatograms from WT, *Rabl2*^*L119F/L119F*^, Rabl2^V158F/V158F^ and *Rabl2*^*-/-*^ mice. The relevant codon bases are labeled above. Black arrow above the chromatogram indicates the insertion. (**c**) qRT-PCR analysis of *Rabl2* expression in testis. Data in **d** are represented as the mean ± SEM and were analyzed using one-way ANOVA with Tukey’s *post hoc* test.

To determine whether these humanized alleles compromised reproduction, we tested fertility by mating, and performed a series of histological and cytological analyses. Whereas female homozygotes had normal litter sizes (Fig 3a), no males homozygous for any of the 3 alleles sired offspring when paired with WT females (Fig 3a). Gross inspection of mutant males revealed phenotypically normal external genitalia, seminal vesicles, and testicular descent. The testis weights from *Rabl2*^*VF/VF*^, *Rabl2*^*LF/LF*^, and *Rabl2*^*-/-*^ mice at 12- and 24-weeks were comparable to age-matched heterozygous and WT littermates (Fig S2a). Histological examination of testis and epididymis sections revealed apparently normal architecture and organized distribution of germ cells, indicating completion of spermatogenesis (Fig S2b and c). As expected, the concentrations of sperm from mutant cauda epididymides were similar to those of heterozygotes and WT (Fig S2d). However, whereas mutant sperm had well-shaped heads and smoothly tapering flagella (Fig S2e), the length of sperm tails were significantly shorter compared to those from heterozygous and WT animals (Fig 3b), consistent with observations of *Rabl2*^*Mot*^ mutant mice ^3^.

**Fig. 3.**
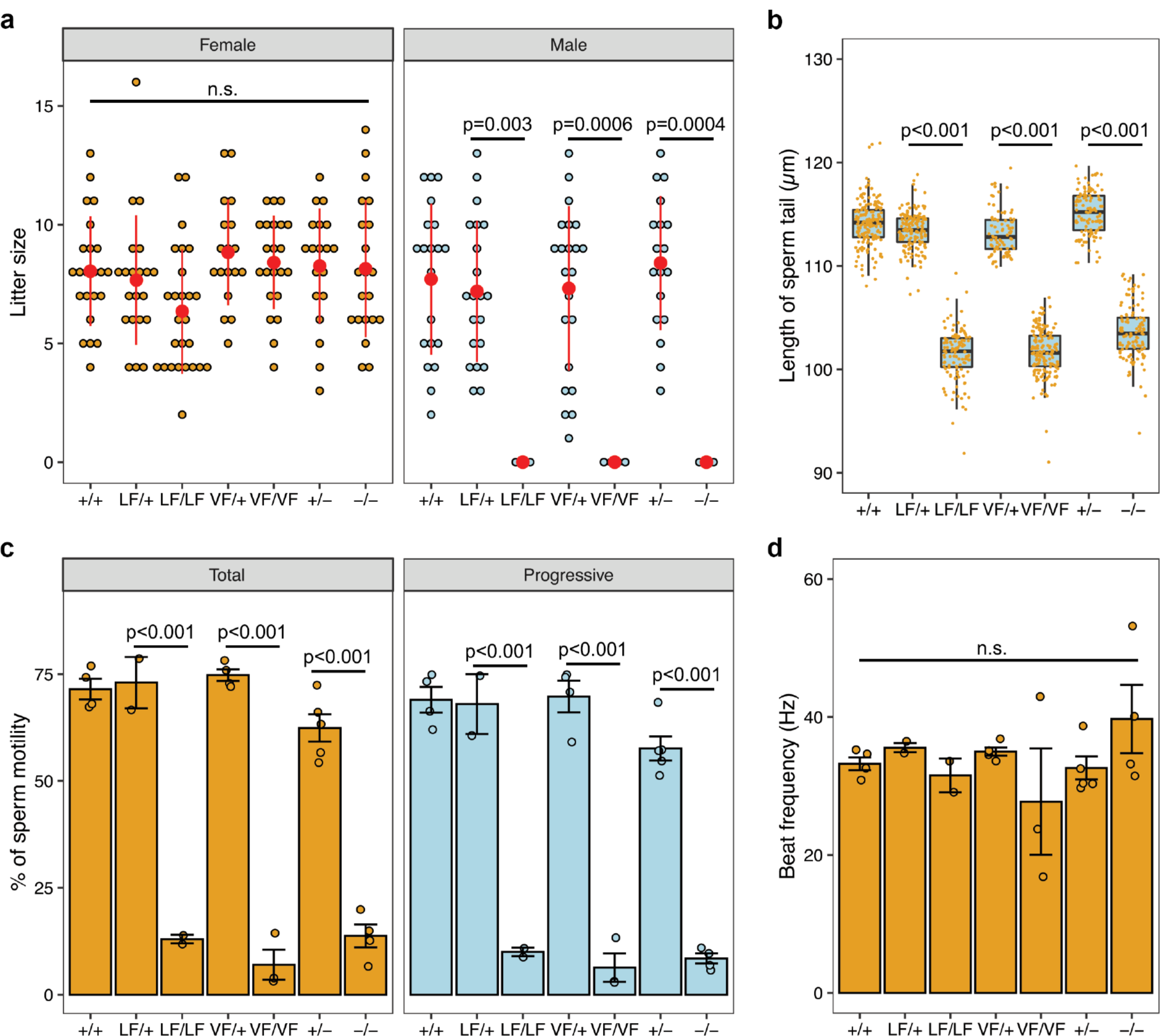
The Rabl2^L119F/L119F^ and Rabl2^V158F/V158F^ alteration cause male infertility. (**a**) Litter sizes from mating of *Rabl2*^*+/+*^ (+/+), *Rabl2*^*L119F/+*^ (LF/+), *Rabl2*^*L119F/L119F*^ (LF/LF), *Rabl2*^*V158F/+*^ (VF/+), *Rabl2*^*V158F/V158F*^ (VF/VF), *Rabl2*^*+/-*^ (+/-) and *Rabl2*^*-/-*^ (-/-) females and males to WT partners. (**b**) Boxplots showing the distribution of the length of sperm tails from +/+ (n=174 from 4 mice), LF/+ (n=158 from 3 mice), LF/LF (n=120 from 3 mice), VF/+ (n=83 from 2 mice), VF/VF (n=160 from 3 mice), +/- (n=120 from 3 mice) and -/- (n=118 from 3 mice) mice. Black central line represents the median, boxes and whiskers represent the 25^th^ and 75^th^, and 2.5^th^ and 97.5^th^ percentiles. (**c**) Assessment of sperm total and progressive motilities of +/+, LF/+, LF/LF, VF/+, VF/VF, +/- and -/- mice. For each mouse, at least 100 sperm were analyzed. (**d**) Beat frequency (in hertz) of motile sperms from +/+, LF/+, LF/LF, VF/+, VF/VF, +/- and -/- mice. Data in **a**-**d** are represented as the mean ± SEM and were analyzed using one-way ANOVA with Tukey’s *post hoc* test. n.s. in **a** and **d** represent no significant difference.

Given that *Rabl2*^*Mot*^ homozygotes are sterile due to defective sperm motility ^3^, we performed CASA (computer assisted sperm analysis) on sperm recovered from the vas deferens of our mutants. *Rabl2*^*LF/LF*^, *Rabl2*^*VF/VF*^ and *Rabl2*^*-/-*^ sperm had significantly reduced motility and progressive movement compared to those from heterozygous and WT males (Fig 3c). However, flagellar beat frequencies were not altered in the few mutant sperm that were motile (Fig 3d), indicating that they were not defective in energy metabolism. Cumulatively, these data suggest that mutant male mice were sterile due to an inability of sperm to reach the oviducts and oocytes in the female genital tract.

Efferent duct motile cilia are necessary to propel immotile sperm from the testis to the epididymis. Loss of efferent duct motile cilia by genetic ablation of miR-34b/c and -449a/b/c causes sperm aggregation and agglutination, as well as luminal obstruction; these defects induce back-pressure atrophy of the testis and ultimately male infertility ^27^. However, histology revealed no indication of efferent duct motile cilia abnormalities in *Rabl2* mutants (Fig. S2f).

### *Rabl2* is important for normal growth, limb development and left-right patterning

Given that *Rabl2* is expressed in many organs ^28^ and its ablation in mouse causes phenotypic hallmarks of PCD and BBS ^3–5^, we extended our observations to nonreproductive systems. Consistent with a previous study ^5^, *Rabl2*^*VF/VF*^, *Rabl2*^*LF/LF*^, *and Rabl2*^*-/-*^ mice became overweight with age (Fig 4a, and Fig S3). However, we noticed that approximately 45% of mutant mice exhibited growth retardation during the lactation period (Fig 4a and Fig S3), despite having neonatal body weights similar to WT and heterozygous siblings. The growth retardation disappeared with age, approximately equalizing by 3 months after birth (Fig 4a). Collectively, these data indicate that *Rabl2* has distinct functions that impact mouse growth.

**Fig. 4.**
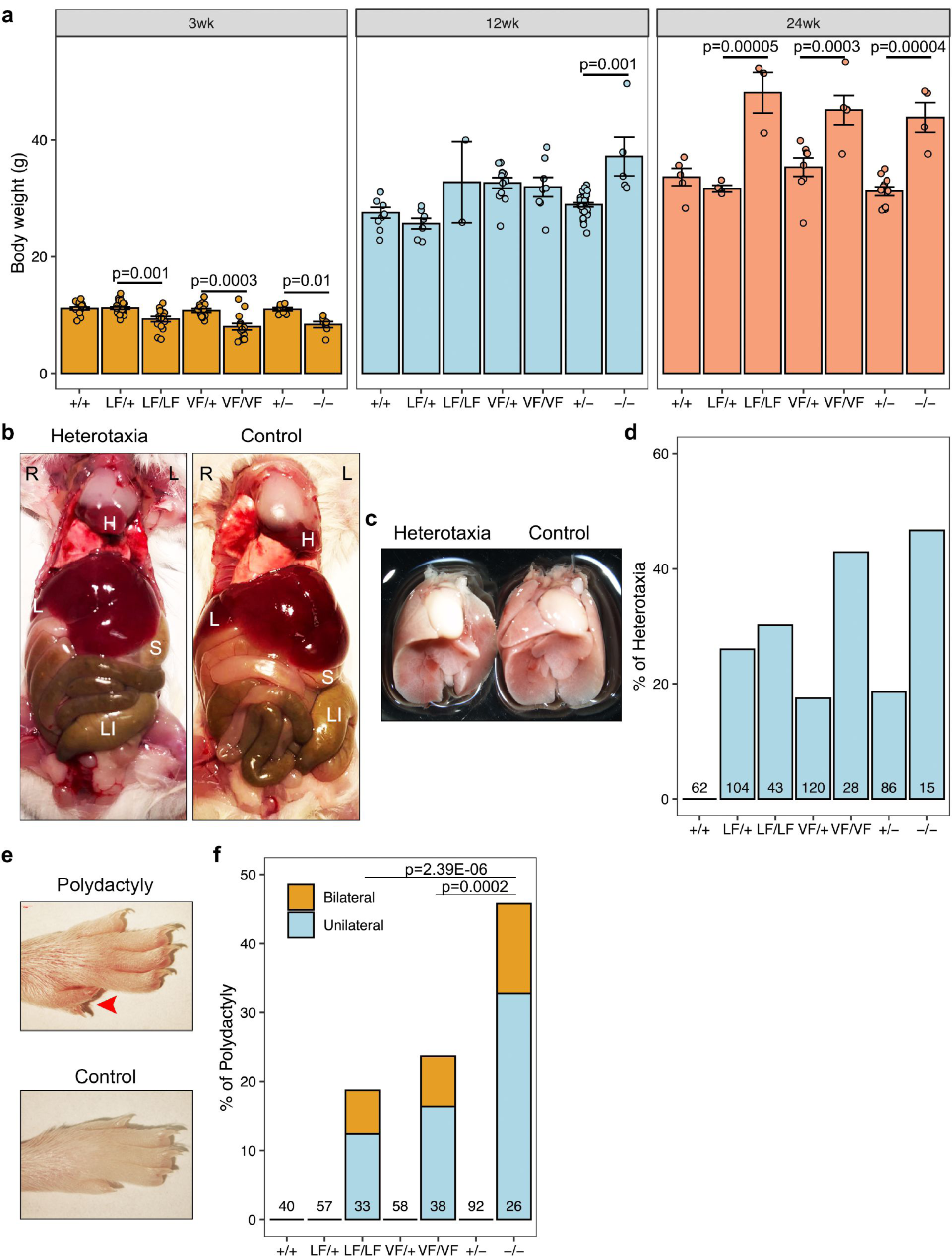
Rabl2 mutant mice have developmental defects. (**a**) Comparison of body weight of +/+, LF/+, LF/LF, VF/+, VF/VF, +/- and -/- males (see legend of Fig. 3 for genotype abbreviations) at 3-, 12- and 24-week (wk) of age. (**b**) Examples of laterality defects in Rabl2 altered mice. Visceral dissection demonstrates the location of heart (H), liver (L), stomach (S) and large intestine (LI). (**c**) Relative location of heart and lung in heterotaxic and control mice. (**d**) Quantification of the proportion of mice with heterotaxia. (**e**) Representative images of the hindlimb from Rabl2 mutant mouse with polydactyly. (**f**) Quantification of the proportion of mice with polydactyly. Data in **a** are represented as the mean ± SEM and were analyzed using one-way ANOVA with Tukey’s *post hoc* test. Data in **f** were analyzed using chi square test.

Inspection of the internal viscera of *Rabl2* mutant mice demonstrated abnormalities in left-right (L/R) axis specification. About 30-50% of homozygous mutants displayed heterotaxia, a partial inversion defect, showing internal organs slightly shifted to the right side (Fig 4b and c). However, we didn’t observe mice with complete situs inversus, a mirror image of normal anatomy. Interestingly, heterotaxia was also found in heterozygotes, albeit at lower frequencies, but not in WT control mice (Fig 4d), suggesting that there may be a certain degree of *Rabl2* haploinsufficiency for certain functions in certain tissues. These findings indicate that a deficiency of RABL2 perturbs mechanisms controlling left-right asymmetry.

Similar to published observations that 67% (6 in 9) of *Rabl2* KO mice had polydactyly ^4^, approximately 19% of *Rabl2*^*LF/LF*^, 24% of *Rabl2*^*VF/VF*^, and 42% of *Rabl2*^*-/-*^ mice exhibited pre-axial polydactyly of the hind feet (Fig 4e and f). In addition, ∼40% of polydactyly appeared on both feet (Fig 4f). Compared with *Rabl2*^*-/-*^, the incidences of polydactyly in *Rabl2*^*VF/VF*^ and *Rabl2*^*LF/LF*^ were lower, suggesting they are hypomorphic alleles for limb development.

### *Rabl2*^*LF/LF*^ and *Rabl2*^*-/-*^ mice have distinct perinatal neural defects

Breeding data from intercrosses of heterozygotes revealed that instead of yielding the expected 25% Mendelian distribution of homozygotes, we found only 14% of *Rabl2*^*LF/LF*^ and *Rabl2*^*-/-*^ live pups at postnatal day 7-14 (Fig 5a), suggesting partially penetrant embryonic lethality. In contrast, *Rabl2*^*VF/VF*^ mice were present at near-Mendelian frequency (23%) (Fig 5a). To identify the stage at which embryonic lethality might be occurring in the former two alleles, we examined embryos at E12.5 and E14.5 from heterozygous intercrosses. We observed embryos with hypoplastic and exencephalic heads (Fig 5b-e), suggesting that *Rabl2* plays a role in neural tube closure. Additionally, we observed an increased incidence of resorption sites at E16.5– E18.5 (Fig 5f). We did not observe liveborn offspring with NTDs, although such pups may have been cannibalized by the mothers immediately after birth.

**Fig. 5.**
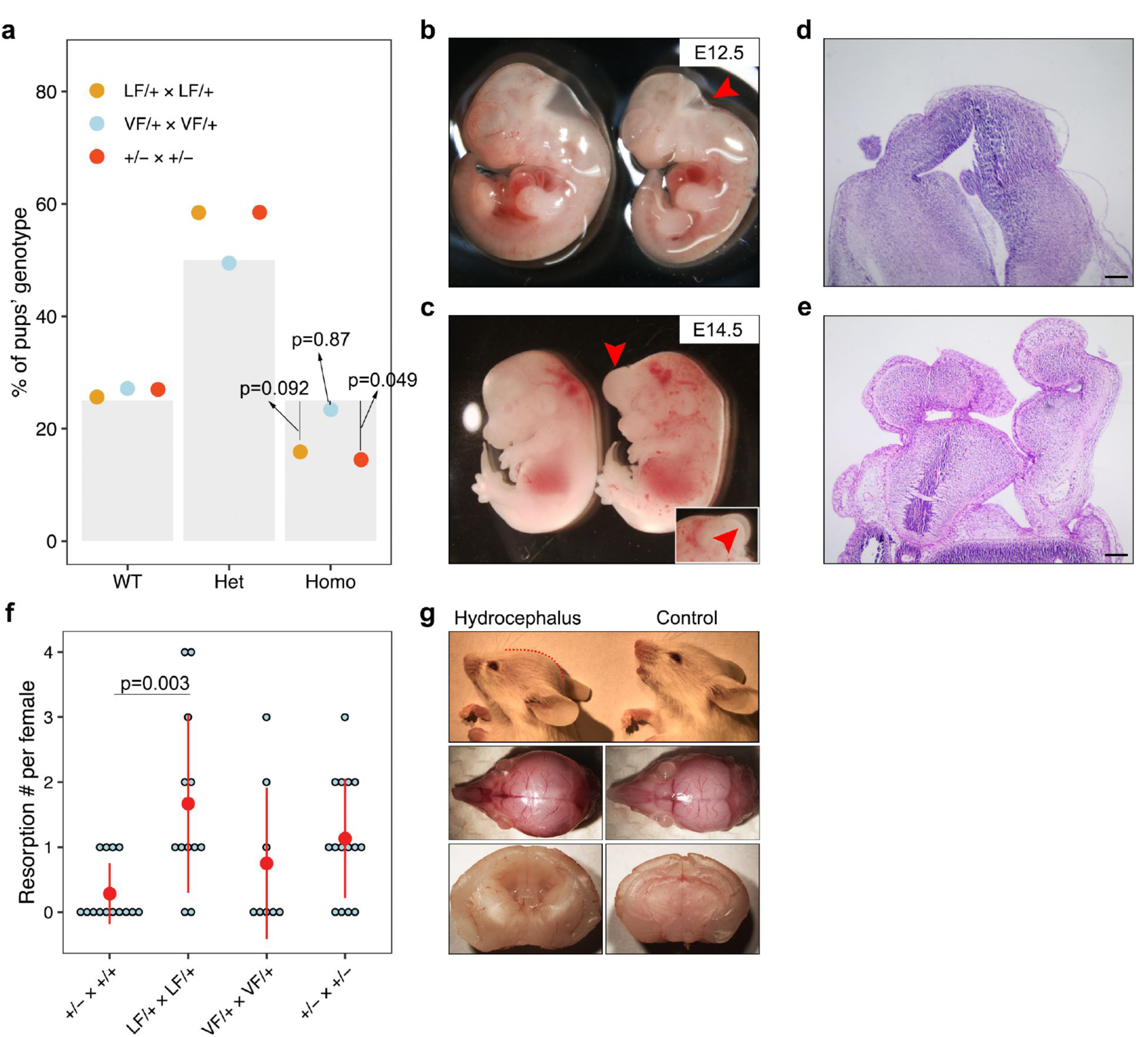
*Rabl2* mutant mice have distinct neural defects. (**a**) Proportion of WT, heterozygous and homologous offspring obtained from het x het intercrosses. The grey bars represent the expected ratios, 25% for WT and homologous and 50% for heterozygous. The dots represent the observed ratios. n=195 offspring obtained from LF/+ intercrosses, n=188 offspring obtained from VF/+ intercrossesand n=200 offspring obtained from +/- intercrosses. (**b**) General observation images of hypoplastic head phenotype (labeled by red arrow) at E12.5 in LF/LF embryo (right panel) as compared with WT control (left panel). (**c**) Exencephaly (labeled by red arrow) in LF/LF (LF = *Rabl2*^*L119F*^) embryo as compared to WT control (left panel) at E14.5. The insertion represents the other side showing the opened head. (**d**) Histology of head for control mouse. (**e**) Cranial histology of mouse with NTD. (**f**) Plot of the number of embryo resorptions per female at E16.5-E18.5 from a heterozygous breeding pair. (**g**) Representative lateral images of 2-wk old LF/LF mouse with a domed cranial vault (red dashed line) compared with WT mouse (upper panel). The middle panel shows the top view of the whole skulls from the same mice. Sagittal sections of brains (lower panel) from a LF/LF and WT mouse revealing dilatation of the lateral and third ventricles in the mutant brain. Data in **a** were analyzed using chi square test. Data in **f** are represented as the mean ± SEM and were analyzed using one-way ANOVA with Tukey’s *post hoc* test. Scale bars in **d** and **e** represent 200 μm.

Finally, we also observed dead or dying pups during the lactation period, partially explaining the lower frequency of homozygous mice observed at the time of genotyping during day 7-14. We occasionally found some *Rabl2* mutant animals with a mildly dome-shaped cranial vault, typical of progressive hydrocephalus (Fig. 5g). Consistent with observations in other another PCD mouse model ^29^, the mice with hydrocephalus always had a smaller body size (Fig. 4a). Collectively, these data suggest that *Rabl2* plays a critical role in the neural tube and cranial development.

## DISCUSSION

A major challenge in medical genetics is to interpret the numerous VUSs in terms of biological and clinical significance. Here, we identified two pathogenic variants in *RABL2A*, rs80006029 and rs200121688, by our integrated computational and experimental approach. Both variants are predicted to be highly deleterious as per the consensus results of SIFT and PolyPhen-2 algorithms. Indeed, *in silico* analysis and *in vitro* protein stability assays found both of alleles destabilized the tertiary structure of RABL2A. These observations, along with the heterogeneous disorders caused by *Rabl2* mutants ^3,5,30^, led us to build humanized mouse models to explore their pathogenesis *in vivo*. Both humanized mice exhibited PCD and BBS features.

*Rabl2* encodes an evolutionarily conserved member of the Rab-like Ras GTPase superfamily. The human paralogs *RABL2A* and *RABL2B* encode nearly identical protein sequences and are expressed in a wide range of tissues, raising the question as to why, or if, both copies are essential. There are three outcomes in the evolution of duplicated genes: (i) one copy may simply become silenced by degenerative mutation; (ii) one copy may acquire a novel, beneficial function and become preserved by natural selection, with the other copy retaining the original function; and (iii) both copies may become partially compromised by mutation accumulation to the point at which their total capacity is reduced to the level of the single-copy ancestral gene ^31^. There are four conserved functional domains within RABL2s: two phosphate-Mg^2+^ binding motifs (G1 and G3), one effector region (G2), and one guanine base-binding motif (G4) ^28^. The effector region determines the specificity of the RAB molecules. In mouse, RABL2 mediates the delivery of effector proteins, including key members of the glycolytic pathway, into the growing sperm tail ^3^. In humans, the effector sequences that differ between RABL2A (DG<u>K</u>TILVDF) and RABL2B (DG<u>R</u>TILVDF) may confer slightly different functions ^28^. Future research will have to address the question of whether human RABL2A and RABL2B have identical or distinct functions during development, spermiogenesis, and/or adult physiology.

We constructed two mouse strains harboring VUS, *Rabl2*^*LF/LF*^ and *Rabl2*^*VF/VF*^, for the primary purpose of determining if they affect fertility. Homozygous null and *Rabl2*^*Mot*^ male mice are infertile ^3,4^, and a deletion allele in human *RABL2A* gene was identified as a risk factor for infertility ^7^. Here, we found that *Rabl2*^*LF/LF*^ and *Rabl2*^*VF/VF*^ mice were infertile, and displayed sperm phenotypes consistent with those described in *Rabl2*^*Mot/Mot*^ mice ^3^. Electron microscopy analysis of *Mot* sperm revealed ultra-structurally normal flagella ^3^. Consistent with this, our CASA data showed that the flagellar beat frequencies in motile *Rabl2*^*LF/LF*^ and *Rabl2*^*VF/VF*^ sperm were comparable to WT and heterozygous controls. The sperm tail truncation is consistent with a role for RABL2 in flagellar assembly, as demonstrated previously ^3^. Given that motile cilia and sperm flagella share the same highly conserved 9+2 microtubule doublet axoneme, and the efferent duct motile cilia are necessary for agitating the sperm to prevent aggregation and clogging during the transition from the rete testis to epididymis ^32^, we checked efferent ductules. However, no defect such as sperm aggregation or agglutination was found in homozygous mutants despite *Rabl2* being expressed in efferent duct motile cilia (https://www.mousephenotype.org/). We propose that efferent duct motile cilia have mild defects and their agitation are also being affected in our *Rabl2* mutant mouse model. However, it is not enough to impede the dominant effect of testicular hydrostatic pressure during the sperm transition.

This study is a part of a larger-scale project to identify human infertility alleles segregating in human populations. However, since *Rabl2* disruption contributes to heterogeneous multisystem disorders such as PCD and BBS ^3,4^ that are caused by cilia defects, we explored the effects of our mutant alleles on nonreproductive systems. Consistent with previous observations ^5^, we found that *Rabl2* mutants become overweight with advancing age (24 wk old). Strikingly, ∼45% exhibited marked growth retardation prior to weaning (3 wks old) before reaching (or in some cases surpassing) the weight of controls by 12 weeks. There are several possible causes of the early growth retardation. One is hydrocephalus, as observed previously in PCD mouse models ^29^. Other potential causes to be investigated are impaired suckling ^32^ or defective release of growth hormone ^33^. The transition to becoming obese in later adulthood suggests a distinct function of RABL2. BBS patients have a high incidence of obesity and diabetes ^34^. A recent study found that elimination of pancreatic β-cell primary cilia diminishes disrupts insulin levels and glucose homeostasis, leading to diabetes ^35^.

Heterotaxia is a birth defect involving randomization of L/R body axis, commonly observed in PCD patients. The leftward-fluid flow generated by embryonic nodal cilia guides the asymmetric morphogenesis of developing organs, and if disrupted, impacts organ lateralization ^36^. Interestingly, we found that not only were *Rabl2* homozygous mutants prone to heterotaxia, but also heterozygotes, indicating that proper RABL2 levels in embryonic nodal cilia are important for establishing proper L/R asymmetry during gastrulation. This, and the commonality of spermiogenesis defects in PCD patients/mouse mutants is consistent with the suggestion that motility functions of cilia (such as that involved in L/R asymmetry) are tightly regulated, and deficiencies in function or quantity are more likely to cause defects related to these functions ^37^. Consistent with this, hydrocephalus is rarer in *Rabl2* mutants than heterotaxia.

Given the abundant expression of *Rabl2* in the brain ^28^, it is likely most if not all the developmental phenotypes of *Rabl2* mutants are a consequence of ciliary defects. The perinatal and juvenile phenotypes such as decreased weight may have a neurological basis, for example, by affecting suckling behavior. NTDs are occasionally present in BBS patients ^38^, and the severe ciliopathies, MKS and JBTS ^39^, commonly display CNS defects. Primary cilia are important for several pathways involved in neural tube patterning, including sonic hedgehog (Shh), wingless-type integration site family (Wnt), and planar cell polarity (PCP). Cerebrospinal fluid (CSF) flow is created by ependymal cells, and is crucial for neurodevelopment and homeostasis of the ventricular system of the brain. The hydrocephalus in *Rabl2* mutant mice may result from an accumulation of CSF due to the decreased ependymal cilia motility. The severity of hydrocephalus in PCD mouse models is affected by inbred strain background, with C57BL/6J mice being particularly severe ^40^. We used a mixed genetic background in these studies (FVB/NJ and B6(Cg)-Tyr^c-2J^/J), possibly explaining the rare incidence of hydrocephalus in our mutants.

In summary, we identified 2 variants in *RABL2A* causing male infertility and ciliopathy by functionally interrogation in humanized mouse models. These genetic variants provide insight into the aetiology of human asthenospermia, PCD and BBS, and can inform clinical management and genetic counseling of patients presenting relevant phenotypes and whom are identified with these variants.

## Acknowledgements

The authors would like to thank R. Munroe and C. Abratte of Cornell’s transgenic facility for generating the edited mice. This work was supported by a grant from the National Institutes of Health (R01 HD082568 to JCS) and contract CO29155 from the NY State Stem Cell Program (NYSTEM). X.D. is supported by a postdoctoral fellowship from the Empire State Stem Cell Fund through New York State Department of Health contract no. C30293GG.

