## Supplemental Figs and Table for "Variants in *RABL2A* causing male infertility and ciliopathy"

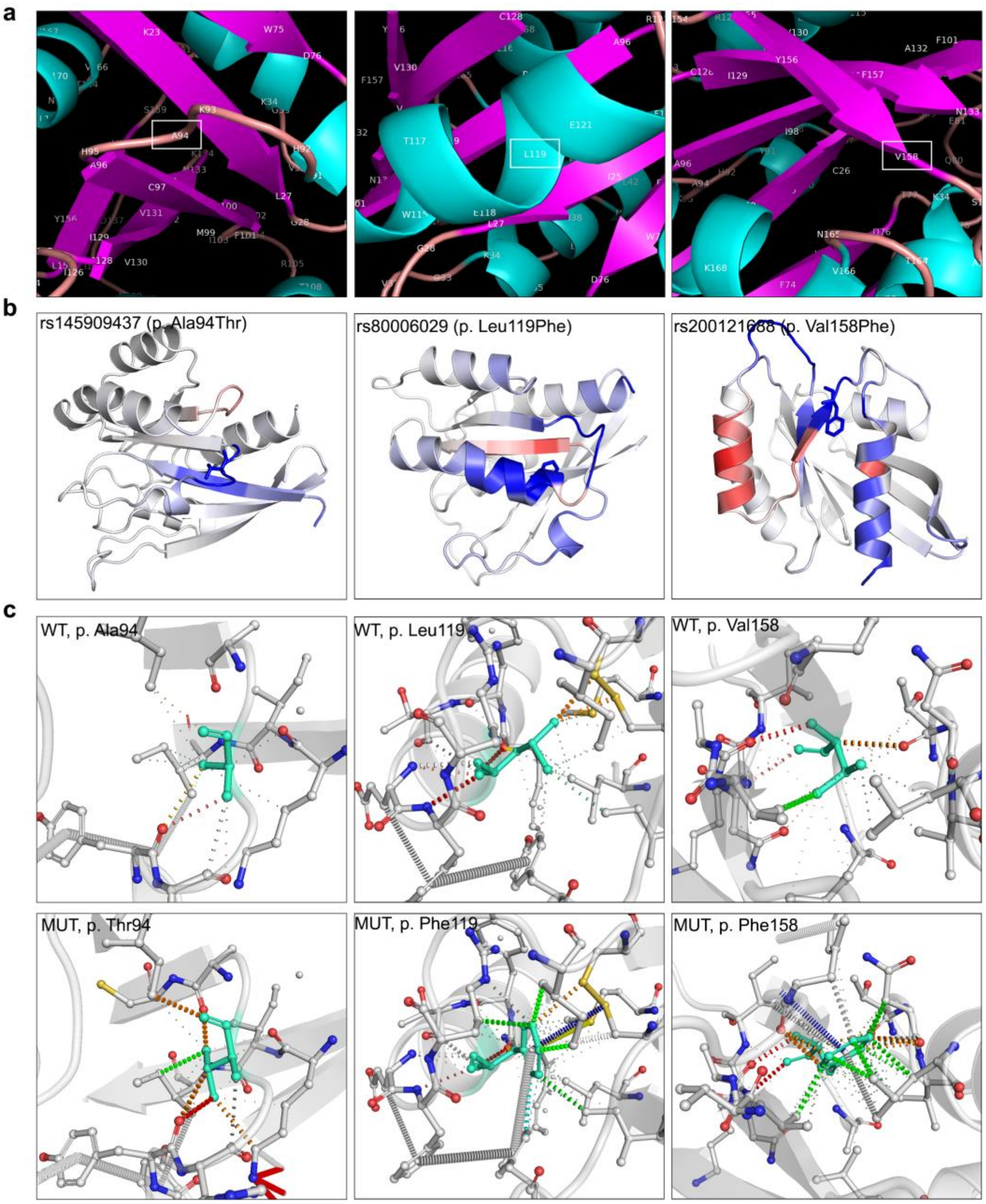


**Fig. S1 Analysis of variants affecting RABL2A protein stability.** (**a**) Close-up view of human RABL2A Ala94, Leu119 and Val158 are located on loop, α-helix and β-sheet, respectively. α-helix, β-sheet and the loop are labeled by different colors. (**b**) Changes in vibrational entropy energy between WT and mutants. Amino acids are colored according to the vibrational entropy change upon mutation. Blue represents a rigidification of the structure and red a gain in flexibility. (**c**) Analysis of contact mediated by residues around mutation sites. WT and mutant residues are colored in light green and are also represented as sticks alongside of the surrounding residues that are involved in any type of interactions. H-bonds are shown in red, VdW (Van der Waals) contacts in gray, hydrophobic-VdW clashes in green, and polar-VdW clashes in orange.

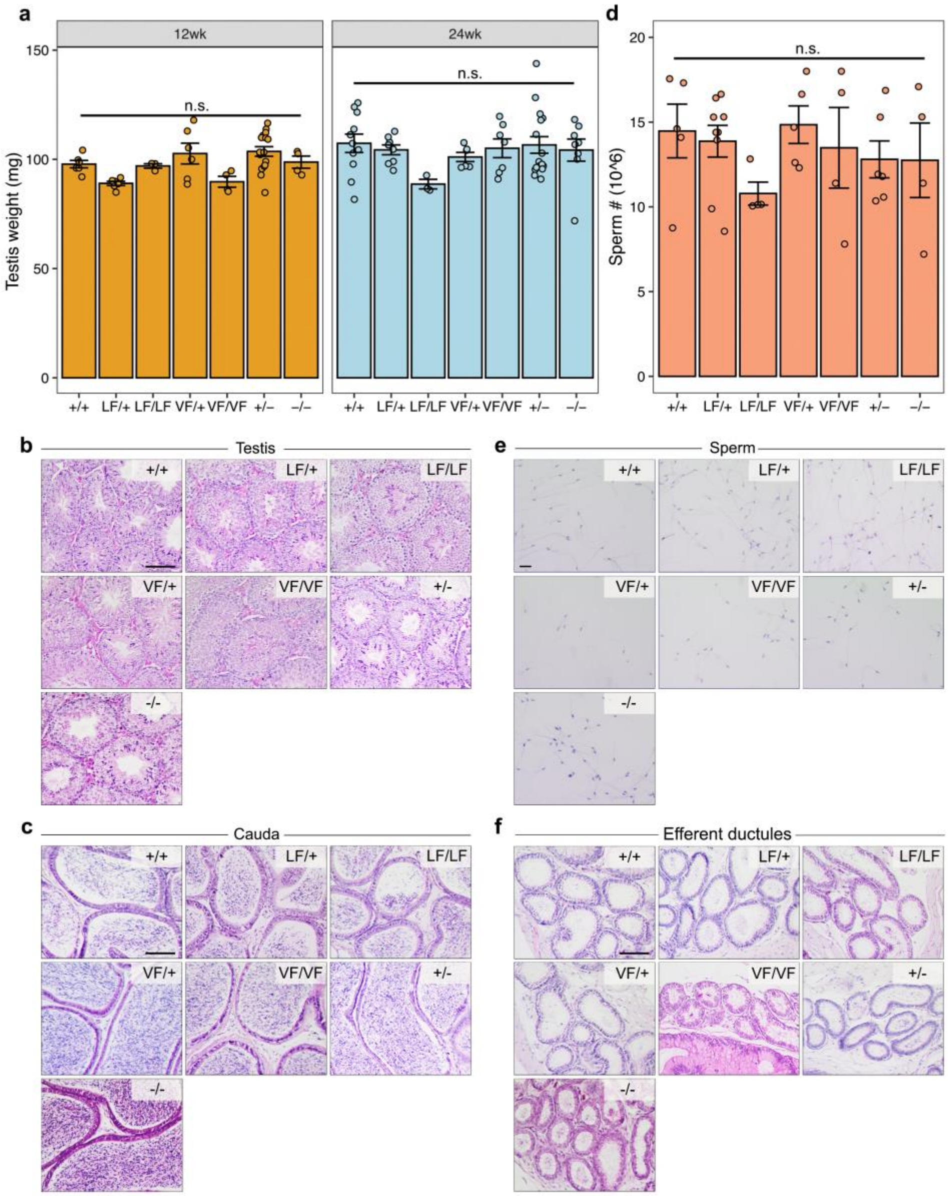


**Fig. S2 Rabl2 mutant mice have no gross abnormalities in testis, epididymis, sperm and efferent ductules.** (**a**) Comparison of testis weights from +/+, LF/+, LF/LF, VF/+, VF/VF, +/- and -/- mice at 12- and 24-wk of age. (**b**) Cross sections of seminiferous tubules from 12-wk old +/+, LF/+, LF/LF, VF/+, VF/VF, +/- and -/- mice. (**c**) Cross sections of cauda epididymis from 12-wk old +/+, LF/+, LF/LF, VF/+, VF/VF, +/- and -/- mice. (**d**) Comparison of sperm number in single cauda epididymis from +/+, LF/+, LF/LF, VF/+, VF/VF, +/- and -/- males. (**e**) H&E stained sperms extracted from the cauda epididymis of +/+, LF/+, LF/LF, VF/+, VF/VF, +/- and -/- males. (**f**) Cross sections of efferent ductules from 24-wk old +/+, LF/+, LF/LF, VF/+, VF/VF, +/- and -/- mice. Data in **a** and **d** are represented as the mean ± SEM and were analyzed using one-way ANOVA with Tukey's *post hoc* test. n.s., no significant difference. Scale bars in **b**, **c** and **f** represent 100 μm, and in **e** represents 20 μm.

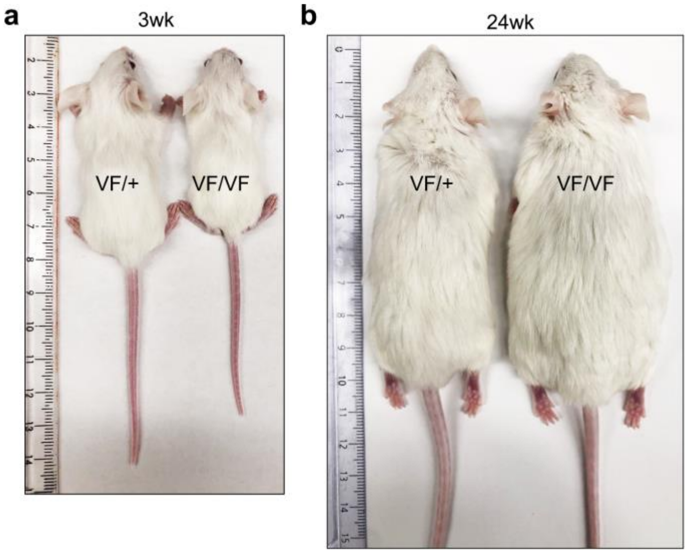


**Fig. S3 Rabl2 mutant mice have growth defects.** (**a**) Representative image of VF/+ and VF/VF mice at 3-wk old. (**b**) Representative image of VF/+ and VF/VF mice at 24-wk old.

Table S1. gRNAs, ssODN and primers sequence used in this study.

| **gRNA/ssODN/Primers** | **Sequence (5′ to 3′)** |
|---|---|
| Rabl2 LF gRNA | ATGCAGAACTTCGGGAGTTC |
| Rabl2 VF gRNA | CAGCAGCCGAGACAAAGTAC |
| Rabl2 LF ssODN | GTATTTGATGTCCAGAGGAAAATCACCTATAAGAACCTGGGTACCTGGTATGCAGAATTT<br>CGGGAGTTTAGACCGGAGATCCCATGCATCCTGGTGGCCAATAAAATTGATGGTGGGGCC |
| Rabl2 VF ssODN | GACTCAAAAGAATTTCAGTTTTGCCAAGAAGTTCTCCCTGCCACTCTATTTTTTCTCGGC<br>TGCTGATGGTACCAATGTTGTGAAGGTAATGGCAGCCCTGAGGCAGTCTGCTGACACCTG |
| Rabl2 LF&KO geno F | CATCTTGGGAAGGGAAACAA |
| Rabl2 LF&KO geno R | TCTTTGCTGTCTGGGACTGA |
| Rabl2 VF geno F | CCCAGCTTTATAACCAGCAG |
| Rabl2 VF geno R | TCCCAAGACACAGACCCATA |
| Rabl2 qPCR F | GATGGTACCAATGTTGTGAAGC |
| Rabl2 qPCR R | AAAACCTCGTCCATGAAGTCC |
| Gapdh qPCR F | CTTTGTCAAGCTCATTTCCTGG |
| Gapdh qPCR R | TCTTGCTCAGTGTCCTTGC |
